# TPH2 knockout male rats are aggressive, show less anxiety, and exhibit an altered oxytocin system

**DOI:** 10.1101/2022.07.08.499134

**Authors:** Xianzong Meng, Joanes Grandjean, Giulia Sbrini, Pieter Schipper, Nita Hofwijks, Francesca Calabrese, Judith Homberg

**Affiliations:** Department of Cognitive Neuroscience, Donders Institute for Brain, Cognition, and Behaviour, Radboud University Medical Centre, Nijmegen, The Netherlands; Department of Pharmacological and Biomolecular Sciences, Università Degli Studi Di Milano, Via Balzaretti 9, 20133, Milan, Italy; Department of Medical Imaging, Radboud University Medical Centre, Nijmegen, The Netherlands

**Keywords:** Tryptophan hydroxylase 2, rats, anxiety, social behavior, oxytocin

## Abstract

The central serotoninergic system is critical for stress responsivity and social behavior, and its dysregulations has been centrally implicated in virtually all neuropsychiatric disorders. Genetic serotonin depletion animal models could provide a tool to elucidate the causes and mechanisms of diseases and to develop new treatment approaches. Previously mice lacking tryptophan hydroxylase 2 (Tph2) have been developed, showing altered behaviors and neurotransmission. However, the effect of congenital serotonin deficiency on emotional and social behavior in rats is still largely unknown, as are the underlying mechanisms. In this study, we used Tph2 knockout (Tph2^-/-^) male rat model to study how the lack of serotonin in the rat brain affects anxiety-like and social behaviors. Since oxytocin is centrally implicated in these behaviors, we furthermore explored whether effects of Tph2 knockout on behavior would relate to changes in the oxytocin system. We show that Tph2^-/-^ rats display reduced anxiety-like behavior and a high level of aggression in social interactions. In addition, oxytocin receptor expression was increased in the infralimbic and prelimbic cortex, paraventricular nucleus, dorsal raphe nucleus and some subregions of hippocampus, which was paralleled by increased levels of oxytocin in the medial frontal cortex, paraventricular nucleus, but not the dorsal raphe nucleus, central amygdala and hippocampus. In conclusion, our study demonstrated reduced anxiety but exaggerated aggression in Tph2^-/-^ male rats and reveals for the first time a potential involvement of altered oxytocin system function.

**SIGNIFICANCE STATEMENT:** We explored the changes in behavior and oxytocin system functioning in the tryptophan hydroxylase 2 (Tph2) knockout rat model, lacking serotonin in the brain. This rat model contributes to our understanding of the role of serotonin in psychiatric transdiagnostic features and underlying mechanisms. We found that Tph2 knockout male rats are aggressive, less anxious, and exhibit an altered oxytocin system. The observed changes in oxytocin signaling may lead to a new target for the treatment of diseases caused by genetic serotonin deficiency.

## Introduction

Serotonin (5-HT) has been long recognized to modulate the stress response and social behavior, and its dysfunction has been implicated in numerous psychiatric disorders. 5-HT synthesis is dependent on the rate-limiting enzyme tryptophan hydroxylase (Tph). There are two Tph isoforms, of which Tph2 is predominantly expressed in the brain (Walther et al., 2003). Indeed, Tph2 mRNA has been detected in multiple brain regions including frontal cortex, thalamus, hippocampus, hypothalamus and amygdala (Zill, Büttner, Eisenmenger, Bondy, & Ackenheil, 2004). The discovery of Tph2 opened up a new area of research. Human studies reported an association between functional Tph2 variants and personality traits (L. Gutknecht et al., 2007) as well as various neuropsychiatric disorders (Waider, Araragi, Gutknecht, & Lesch, 2011).

Animals with targeted deletion of genes encoding mediators of the serotonergic transmission have been proven to be a powerful tool for detailed understanding contributions of the genetic basis of traits related to mood disorders. To model human Tph2 gene variance, Tph2 knockout (Tph2^-/-^) mice have been generated. Although they do not exactly mimic human Tph2 polymorphisms, the animals show phenotypes that are grossly in line with the humane gene-association studies. More specifically, Tph2^-/-^ males show more aggression (Angoa-Pérez et al., 2012; Mosienko et al., 2012). Even female Tph2^-/-^ mice and weanlings (3-4 weeks old) of both sexes showed elevated aggressive in a modified resident-intruder test (Angoa-Pérez et al., 2012). Furthermore, increased obsessive-compulsive-like behavior was observed in Tph2^-/-^ mice in the marble burying test (Angoa-Pérez et al., 2012; Savelieva et al., 2008). Tph2^-/-^ mice show no difference in total locomotor activity or exploratory behaviors in the open-field test, but they spent less time in the central field, indicative for elevated anxiety-like traits (Savelieva et al., 2008). In some studies it is also reported that Tph2^-/-^ mice either displayed marginally reduced anxiety- and depression-like behavior (Lise Gutknecht et al., 2015), or do not display a depression-like behavioral phenotype (Angoa-Pérez et al., 2014).

Tph2^-/-^ rats were introduced in 2016 (Kaplan et al., 2016). Studies employing Tph2^-/-^ rats showed increased aggressive behavior (Peeters et al., 2019), and increased neuroplasticity in basal condition (Brivio et al., 2018), and an impaired response to acute stress exposure (Brivio et al., 2018; Sbrini, Brivio, Bosch, Homberg, & Calabrese, 2020). However, at the behavioral level the study of Tph2^-/-^ rats is still inadequate. As to whether the rat model also demonstrates anxiety- and depression-like phenotypes and further social disturbances like in Tph2^-/-^ mice remains to be established, as well as the potential underlying neurobiological mechanisms.

Taking human and mouse Tph2 data together, the changes in the expression of enzyme appears to particularly affect the domains of affective and social behavior. One molecule that is centrally implicated in both these behavioral domains is oxytocin. In animal studies, oxytocin was firstly indicated being involved in depressive behaviors originated from the finding that intracerebroventricular oxytocin administration diminished the immobility time in mice in the forced swimming test (Meisenberg, 1981). After that, it has been shown that intraperitoneal oxytocin administration reduced the immobility in this test (Arletti & Bertolini, 1987). The role of oxytocin in depression-related behavior was getting increasing attention due to the findings that this hormone plays an important role in social attraction, affiliative behavior and bonding, which could be potentially important in relation to the development of depression (Insel & Young, 2001; Neumann, 2008).

Because 5-HT and oxytocin both have effects on anxiety and social processes, the attention for interactions between 5-HT and oxytocin is increasing. Central administration of selective 5-HT agonists increased the expression of oxytocin mRNA in hypothalamic nuclei (Jørgensen, Kjær, Knigge, Møller, & Warberg, 2003), which is consistent with reports that 5-HT and 5-HT fibers influence brain regions rich in oxytocin (Emiliano, Cruz, Pannoni, & Fudge, 2007; Ho, Chow, & Yung, 2007; Sawchenko, Swanson, Steinbusch, & Verhofstad, 1983). Central injection of oxytocin reduces anxiety in the rat social interaction test, which is fully blocked by an antagonist of 5-HT2A/2C receptors (Yoshida et al., 2009).

Based on the above, we hypothesized that the behavioral characteristics of Tph2^-/-^ rats is related to altered oxytocin signaling. To test this hypothesis, we determined oxytocin levels in brain regions that have been reported to be mediated by oxytocinergic mechanisms effecting social and aggressive behaviors as well as expression levels of oxytocin receptors which play a key role in these traits.

## Materials and methods

### Animals

Tph2 knockout (Tph2^-/-^) rats were generated by a truncation mutation (Hodges, Kaplan, Echert, Puissant, & Geurts, 2015). Tph2^-/-^, wild-type (Tph2^+/+^) and heterozygous (Tph2^+/-^) rats were derived by crossing heterozygous rats (dark agouti) that were out crossed with wild-type rats (DA/OlaHsd) (Jacob Human and Molecular Genetics Center, Medical College of Wisconsin, Milwaukee, USA). For behavioral testing, twenty-six male rats (n_Tph2+/+_ = 10, n_Tph2-/-_ = 7, n_Tph2+/-_ = 9) were housed 2-3 per cage (25 × 25 × 35 cm3, length x width x height) with 2 cm sawdust bedding in a 12 h light-dark cycle from 8 am to 8 pm at a temperature of 21±1°C under controlled environmental conditions (humidity 45-60%), with food and water provided ad libitum. Rats between 70 ± 14 days old were used for all experiments, exclusively during the light period. For molecular testing, another cohort of twenty rats (n_Tph2+/+_ = 10, n_Tph2-/-_ = 10) were housed under same conditions. All efforts to retain animals as humane as possible were made according to the three Rs for all animals used (Russell & Burch, 1959).

All procedures were executed in accordance with the Dutch legal ethical guidelines of animal experiments, as approved by the Central Committee Animal Experiments, the Hague, the Netherlands.

### Elevated plus maze

Anxiety-like behavior was measured using the elevated plus maze. The maze, elevated 50 cm from the floor, consisted of two open arms (50 × 10 cm, 10 lux) and two closed arms (50 × 10 cm) that were enclosed by a side wall. Rats were placed in the center of the maze, facing the open arm and could freely explore the apparatus for 5 min (Pellow, Chopin, File, & Briley, 1985), while being recorded by a camera suspended above the center of the maze. Total open and closed arm entries, duration and latency as well as total distance travelled on all arms were quantified. Results were collected using Observer Ethovision version (Noldus, Wageningen, the Netherlands) by a researcher blind to treatment conditions.

### Social behavior

Two unfamiliar animals with the same genotype were exposed to each other in a novel context for 20 min after being isolated for 3.5 h in a separate housing room. The novel context consisted of a Phenotyper cage (45 × 45 × 45 cm3) with standard sawdust bedding (2 cm). Rats had no access to food or water during the experiment. Each 20 min session was recorded, and videos were scored using J-Watcher version 1.0 (Dan Blumstein’s Lab, University of California, Los Angeles; The Animal Behavior Lab, Macquarie University, Sydney, Australia). Social interaction and aggressive interaction parameters for each individual rat were scored by the same experimenter according to Table 1. The data from two Tph2^-/-^ rats were removed from the analysis because of a fierce fight between the two animals, which ended with one of the rats hiding in a corner and not moving anymore.

**Table 1.** Social interaction and aggressive interaction behaviors measured during the social interaction test.

| Social interaction | Aggressive interaction |
| --- | --- |
| No contact | Aggressive behavior |
| Self-grooming | Mounting |
| Rearing | Chasing |
| Social exploration | Defending |
| Grooming the other | Running |
| Total no contact includes no contact, self-grooming and rearing;<br>Total aggressive behaviors includes aggressive behavior, mounting and chasing. |  |

### Analysis of oxytocin receptor mRNA expression levels

To eliminate the effects from behavioral testing on gene expression, another independent group of rats was used for a molecular study for which we used Tph2^+/+^ and Tph2^-/-^ rats. The rats were sacrificed through decapitation and immediately frozen at −80 °C. The left hemisphere was used for qPCR. Brain regions were dissected according to The Rat Brain in Stereotaxic Coordinates 6th Edition (Paxinos & Watson, 2006) by brain punching using a Cryostat machine. We punched out the prelimbic cortex (Bregma 4.20mm ~ 2.52mm), infralimbic cortex (Bregma 3.72mm ~ 2.52mm), paraventricular thalamic nucleus (Bregma −1.20mm ~ −3.96mm), central amygdaloid nucleus (Bregma −1.44mm ~ −3.24mm), granular layer of the dentate gyrus (dorsal) (Bregma −2.16mm ~ −3.00mm), granular layer of the dentate gyrus (ventral) (Bregma −4.36mm ~ −5.04mm), field CA1 of the hippocampus (dorsal) (Bregma −2.52mm ~ −3.00mm), field CA1 of the hippocampus (ventral) (Bregma −4.36mm ~ −5.04mm), field CA3 of the hippocampus (dorsal) (Bregma −2.52mm ~ −3.00mm), field CA3 of the hippocampus (ventral) (Bregma −4.36mm ~ −5.04mm), and the dorsal raphe nucleus (Bregma −6.96mm ~ −8.40mm). The location of the brain punches is shown in Figure 3. Total RNA was isolated by a single step of guanidinium isothiocyanate/phenol extraction by using a PureZol RNA isolation reagent (Bio-Rad Laboratories, Segrate, Italy) according to the manufacturer’s instructions and quantified by spectrophotometric analysis. The samples were then processed for real-time polymerase chain reaction (RT-PCR) to assess the expression of the oxytocin receptor (primers and probe assay ID: Rn00564446_g1, purchased from Life Technologies). In particular, an aliquot of each sample was treated with DNAse (Thermoscientific, Rodano, Italy) to avoid DNA contamination. Purified RNA was analyzed by TaqMan qRT-PCR one-step RT-PCR kit for probes (Bio-Rad laboratories, Italy) with a TaqMan RT-PCR instrument (CFX384 real time system, Bio-Rad Laboratories). After the initial retrotranscription step, 39 cycles of PCR were performed. Samples were run in 384 well formats in triplicate as multiplexed reactions with a normalizing internal control (*36b4;* Forward primer: TTCCCACTGGCTGAAAAGGT; Reverse primer: CGCAGCCGCAAATGC; Probe: AAGGCCTTCCTGGCCGATCCATC, purchased from Eurofins MWG-Operon, Germany). A comparative cycle threshold (Ct) method was used to calculate the relative target gene expression.

### Analysis of oxytocin levels

The right hemisphere was used to measure oxytocin levels. We focused on the medial frontal cortex, paraventricular thalamic nucleus, dorsal raphe nucleus, central nucleus of the amygdala and the hippocampus. Due to the detection range limit, we pooled the CA1, CA3 and dentate gyrus regions from the ventral and dorsal parts of the hippocampus. Brain regions were punched using the same method as described above. Then the brain punching samples were homogenated in RIPA buffer (Sigma, lot. R0278) with Proteinase inhibitor (Thermo Scientific™ Halt™ Protease Inhibitor Cocktail, Lot. WF327612). The location of the brain punches is shown in Figure 3. After centrifugation at in 4°C at 10,000 rcf for 10 min, the supernatant was collected and diluted by PBS. The protein concentration was measured using Micro BCA Protein Assay Kit (ThermoFisher, lot. WF325481). Finally, the supernatant calibrated into the same protein concentration was used for the measurement of oxytocin levels using an ELISA kit (Abcam, lot. 133050), according to the manufacturer’s instructions.

### Statistical analysis

Statistical inference was chiefly based on effect size (Hedges’g) and confidence intervals. P-values were estimated using non-parametric permutation tests. Confidence intervals and p-values were estimated by shuffling the group labels over 5000 permutations. The results are represented as Gardner-Altman plots and reported in the text as effect size [lower bound; upper bound of 95% confidence interval], p value. Effect size interpretations follow Cohen’s 1998 guidelines (Pellow et al., 1985). Small effect: g > 0.2; medium effect: g > 0.4; large effect: g > 0.8. The code and the table to reproduce this analysis are provided freely: https://gitlab.socsci.ru.nl/preclinical-neuroimaging/tph2.

## Results

### Reduced anxiety in Tph2 knockout rats

Elevated plus maze is a classic assay to assess anxiety levels. Tph2^-/-^ rats spent more time in the open arms relative to Tph2^+/+^ rats (Figure 1A, g _Tph2+/+ < Tph2-/-_ = 1.05 [0.17; 2.05], p = 0.04), indicating a lower anxiety level in Tph2^-/-^rats. Consistently, Tph2^-/-^ rats entered closed arms less often compared to Tph2^+/+^ rats (Figure 1B, g _Tph2+/+ < Tph2-/-_ = 1.1 [-2.35; 0.08], p = 0.03). Notably, there is also a medium, albeit non-significant, effect between Tph2^+/+^ and Tph2^+/-^ groups (Figure 1B, g _Tph2+/+ < Tph2+/-_ = 0.56 [-1.43; 0.40], p = 0.22). In other words, the fewer Tph2 gene copies, the less frequent the rats enter closed arms.

**Figure 1.**
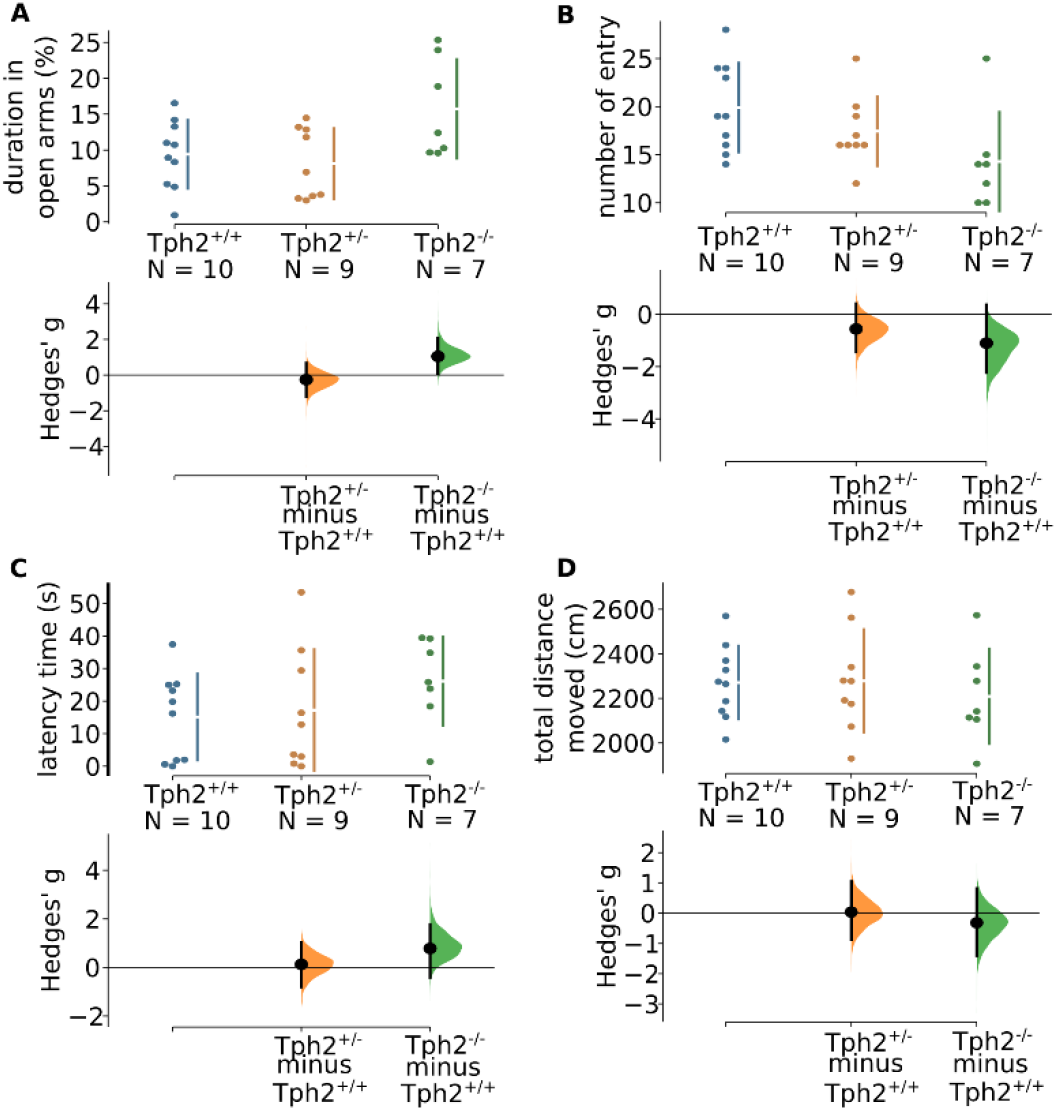
Elevated plus maze test. (A) time spent in open arms, (B) closed arms entries, (C) latency to enter open arms, (D) total distance moved on the elevated plus maze. n_Tph2+/+_ = 10, n_Tph2-/-_ = 7, n_Tph2+/-_ = 9. The Hedges’ g for 2 comparisons against the shared control Tph2^+/+^ are shown in the Cumming estimation plot. The raw data are plotted on the upper axes. On the lower axes, mean differences are plotted as bootstrap sampling distributions. Each mean difference is depicted as a dot. Each 95% confidence interval is indicated by the ends of the vertical error bars.

The latency of the first entry into the open arms is a less conventional anxiety-related parameter but is of interest as it reflects the approach-avoidance conflict concerning aversive open arms. In our experiment, we did not find any noticeable effect between Tph2^+/+^ and Tph2^+/-^ groups (figure 1C). However, a trending effect was found between Tph2^+/+^ and Tph2^-/-^ groups (g _Tph2+/+ < Tph2-/-_ = 0.78 [-0.32; 1.87], p = 0. 12), which suggests Tph2^-/-^ rats have a higher latency of entering into aversive arms. A higher latency is sometimes interpreted as a sign for elevated anxiety level in rodents. However, in our case, taking all the above information into consideration, we interpret this phenomenon as Tph2^-/-^ rats displaying a lower sensitivity to the environment.

Finally, locomotor activity was evaluated by checking total distance rats traveled on elevated plus maze. We could not establish a difference between Tph2^+/+^ and Tph2^+/-^ or between Tph2^+/+^ and Tph2^-/-^ groups (Figure 1D, g _Tph2+/+ < Tph2-/-_ = −0.32 [-1.37; 0.75], p = 0. 51). In conclusion, there is no discernible differences in the locomotor activity among three groups. We therefore conclude that differences in the elevated plus maze assay reflect reduced anxiety levels in the Tph2^-/-^ rats, which is not due to a change in locomotor activity.

### Elevated aggressiveness in Tph2 knockout rats

Following the elevated plus maze test (24 hours later), two unfamiliar rats from the same genotype were exposed to each other in a novel context for 20 min after being isolated for 3.5 h in a separate housing room (Figure 2A). We found a large genotype effect on total no contact behavior (Figure 2B, g _Tph2+/+ < Tph2-/-_ = −6.38 [−7.98; −4.5], p < 0.01), indicating that Tph2^-/-^ rats have a higher level of active social interaction compared with Tph2^+/+^ rats. However, the prolonged social interaction of Tph2^-/-^ rats manifested as increased mounting behaviors. Indeed, Tph2^+/-^ groups showed a trend towards more mounting behaviors than Tph2^+/+^ group (Figure 2C, g _Tph2+/+ < Tph2+/-_ = 0.64 [-0.42; 1.36], p = 0.16). Meanwhile, mounting behavior was significantly increased in Tph2^-/-^ in comparison with Tph2^+/+^ rats (g _Tph2+/+ < Tph2-/-_ = 1.47 [0.80; 3.83], p = 0.01). In other words, the disruption of Tph2 gene leads to more mounting behavior.

**Figure 2.**
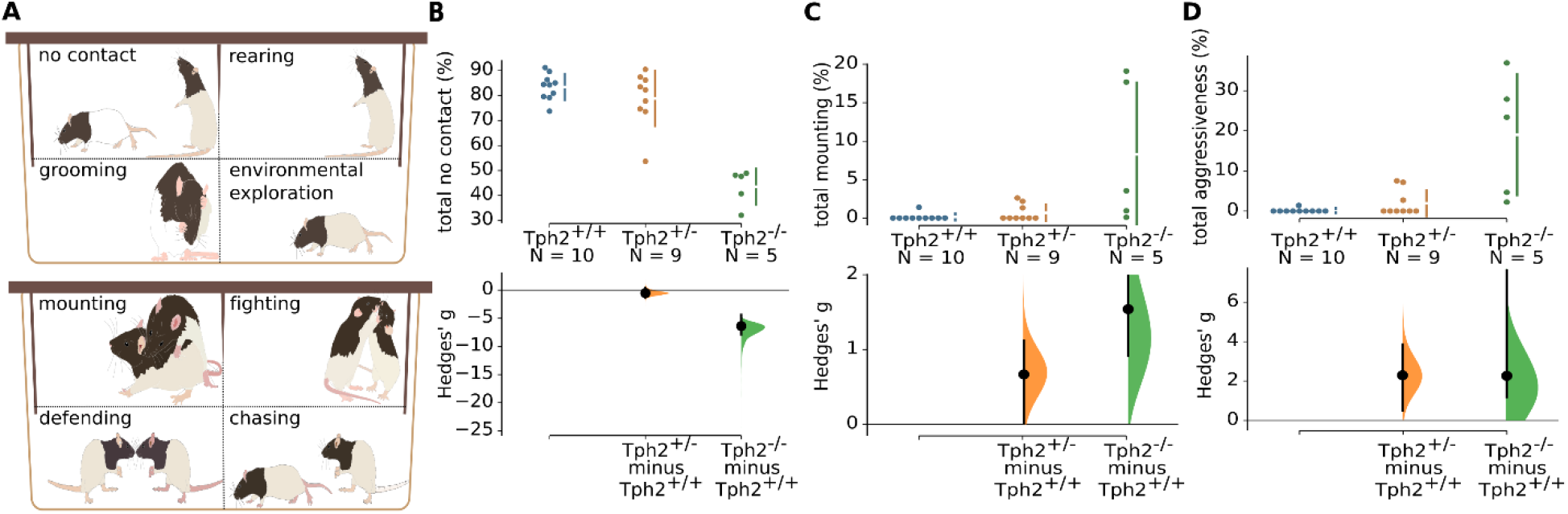
Social behavior test. (A) behavioral categories, (B) total no contact (%), (C) total mounting (%), (D) total aggressiveness (combined time mounting, fighting, defending, and chasing, %). n_Tph2+/+_ = 10, n_Tph2-/-_ = 7, n_Tph2+/-_ = 9. The Hedges’ g for 2 comparisons against the shared control Tph2^+/+^ are shown in the Cumming estimation plot. The raw data are plotted on the upper axes. On the lower axes, mean differences are plotted as bootstrap sampling distributions. Each mean difference is depicted as a dot. Each 95% confidence interval is indicated by the ends of the vertical error bars.

Inter-male mounting may be a marker for dominance or aggressiveness. Finally, we assessed the total time spent on aggressiveness, which included aggressive behaviors, mounting, and chasing behaviors all together. We found that Tph2^-/-^ and Tph2^+/-^ male rats spent more time on aggressive behaviors compared to wild-type controls (Figure 2D, g _Tph2+/+ < Tph2-/-_ = 2.13 [1.09; 7.78], p < 0.01, g _Tph2+/+ < Tph2+/-_ = 0.77 [0.184; 1.56], p = 0.10). We concluded that Tph2 gene knockout is sufficient to increase aggressiveness in male rats.

### Altered oxytocin receptor mRNA expression in Tph2 knockout rats

We found that homozygous and heterozygous Tph2 knockout was sufficient to alter both anxiety and aggressive behaviors in male rats relative to wild-type controls. Due to its role in intensive interactions with serotonin, we proposed that oxytocin may be a relevant mediator. To test this, we first examined oxytocin receptor gene expression (mRNA levels) in areas previously associated with anxiety and aggression (Figure 3A). We presented 4 subregions to parallel the receptor and oxytocin levels in Figure 3, while some other data was presented in Table 2.

**Figure 3.**
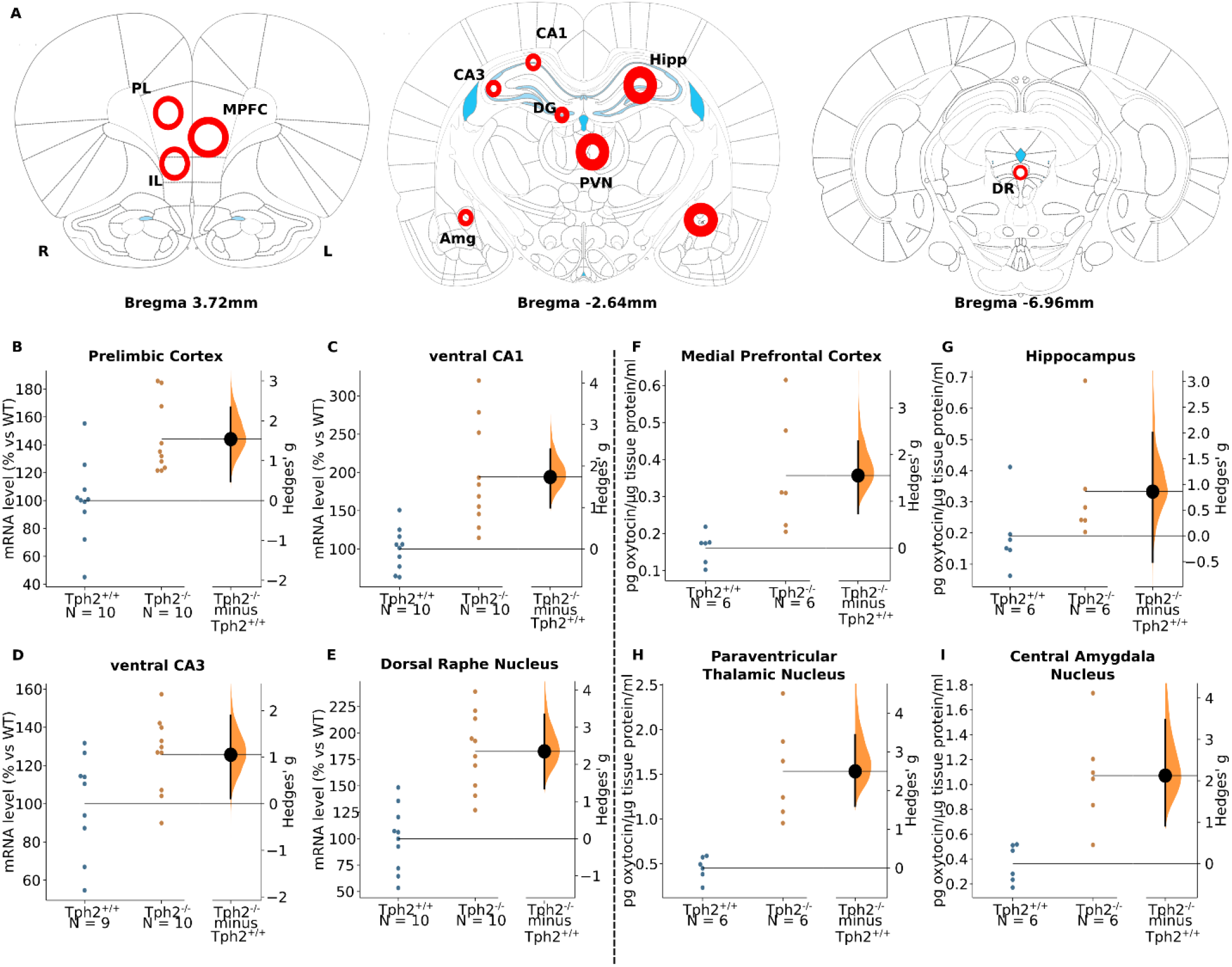
Oxytocin receptor mRNA expression and oxytocin levels (left side: oxytocin receptor mRNA expression; right side: oxytocin level). (A) brain punching sites diagram, (B) prelimbic cortex, (C) ventral CA1 region, (D) ventral CA3 region, (E) dorsal raphe nucleus, (F) medial frontal cortex, (G) hippocampus, (H) paraventricular thalamic nucleus, (I) central nucleus of the amygdala. For oxytocin ELISA results, n = WT (6), Tph2^-/-^ (6), for oxytocin receptor PCR results, _Tph2+/+_ = 10, n_Tph2-/-_ = 10. The Hedges’ g between Tph2^+/+^ and Tph2^-/-^ is shown in the above Gardner-Altman estimation plot. Both groups are plotted on the left axes; the mean difference is plotted on floating axes on the right as a bootstrap sampling distribution. The mean difference is depicted as a dot, the 95% confidence interval is indicated by the ends of the vertical error bar. Abbreviations: PL, prelimbic cortex; MPFC, medial prefrontal cortex; IL, infralimbic cortex; CA1, field CA1 of the hippocampus; CA3, field CA3 of the hippocampus; DG, granular layer of dentate gyrus; PVN, paraventricular thalamic nucleus; Amg, central amygdala nucleus; Hipp, hippocampus; DR, dorsal raphe nucleus.

**Table 2.** The oxytocin receptor mRNA expression levels in different brain regions.

| Location | Hedge's $g$ [95%CI]<br>$Tph2^{+/+} < Tph2^{-/-}$ | P<br>value |
| --- | --- | --- |
| Infralimbic cortex | 1.14 [0.07; 2.18] | 0.02 |
| Dorsal dentate gyrus | 0.93 [-0.02; 1.64] | 0.05 |
| Ventral dentate gyrus | 1.22 [0.32; 2.05] | 0.02 |
| Paraventricular thalamic<br>nucleus | 1.49 [0.52; 2.66] | <0.01 |
| Central nucleus of the<br>amygdala | 0.44 [-0.52; 1.31] | 0.32 |

Oxytocin receptor mRNA expression levels were found to be increased in the infralimbic cortex (Table 2, g _Tph2+/+ < Tph2-/-_ = 1.14 [0.07; 2.18], p = 0.02), paraventricular nucleus (Table 2, g _Tph2+/+ < Tph2-/-_ = 1.49 [0.52; 2.66], p < 0.01), prelimbic cortex (Figure 3B, g _Tph2+/+ < Tph2-/-_ = 1.54 [0.44; 2.33], p < 0.01), and dorsal raphe nucleus (Figure 3E, g _Tph2+/+ < Tph2-/-_ = 2.35 [1.34; 3.37], p < 0 .01). In this study, the hippocampus was functionally segmented into dorsal and ventral compartments, and three regions were tested including CA1, CA3 and granular layer of dentate gyrus. In the dorsal hippocampal compartment, the expression of oxytocin receptors was largely increased in dentate gyrus (Table 2, g _Tph2+/+ < Tph2-/-_ = 0.93 [-0.02; 1.64], p = 0.05). In the CA3 region a small change was found, and no change was found in the CA1 region.

However, in the ventral hippocampal compartment, the expression in CA1 (Figure 3C, g _Tph2+/+ < Tph2-/-_ = 1.74 [0.99; 2.39], p < 0.01), CA3 (Figure 3D, g _Tph2+/+ < Tph2-/-_ = 1.05 [0.13; 1.91], p = 0.03) and dentate gyrus (Table 2, g _Tph2+/+ < Tph2-/-_ = 1.22 [0.32; 2.05], p = 0.02) were all largely increased. We conclude that oxytocin receptor expression was elevated consistently throughout the brain in Tph2^-/-^ relative to Tph2^+/+^ rats.

### Altered oxytocin levels in Tph2 knockout rats

In addition to examining receptor expression levels, we also determined oxytocin concentration (Figure 3A). Because of the sensitivity of the assay, several areas were merged to achieve sufficient peptide levels (e.g., prelimbic and infralimbic cortex). This is justified because of the indiscriminate receptor mRNA elevation in the pooled regions. Our oxytocin ELISA results indicated that the oxytocin level was largely increased in the medial prefrontal cortex (Figure 3F, g _Tph2+/+ < Tph2-/-_ = 1.55 [0.80; 2.3], p = 0.02), hippocampus (Figure 3G, g _Tph2+/+ < Tph2-/-_ = 0.86 [-0.64; 1.94], p = 0.14), paraventricular thalamic nucleus (Figure 3H, g _Tph2+/+ < Tph2-/-_ = 2.5 [1.69; 3.63], p < 0 .01) and central nucleus of the amygdala (Figure 3I, g _Tph2+/+ < Tph2-/-_ = 2.13 [0.9; 3.57], p < 0.01). We conclude that, similar to the oxytocin receptor, the ligand is found more abundantly in the areas sampled of Thp2^-/-^ male rats, relative to wild-type controls.

## Discussion

The results from this study reveal that the knockout of Tph2 significantly affects rat’s behavior and influences oxytocin levels and the expression of its receptors. Tph2^-/-^ rats are less anxious and show more social interaction. However, social interaction is dominated by high levels of aggression and mounting.

Tph2^-/-^ rats exhibited less anxiety-like behaviors in the elevated plus maze as supported by a longer duration in open arms and a reduction in closed arms entries. However, if the rats were less anxious, rats should have entered the open arm quicker but data we collected showed the opposite. Contrary to Tph2^-/-^rats, serotonin transporter knockout rats, which harbor a high brain serotonin concentration, showed high sensitivity to environmental stimuli (Homberg & Lesch, 2011; Sbrini et al., 2020). Hence, it is possible that the reduced anxiety level of Tph2^-/-^ rats relates to an attenuated environmental sensitivity, reducing awareness of the difference between the open and closed arms. At the same time, the decreased anxiety level is independent of activity, as total distance traveled does not differ between genotypes. Interestingly, an 82% serotonergic neurotoxin-induced depletion of 5-HT in the rat medial prefrontal cortex increased anxiety-like behavior on the elevated plus-maze (Pum, Huston, & Müller, 2009). Given the fact that the depletion of serotonin *ab origine* probably leads to compensatory responses as often seen in conventional knockout animal model (Knobelman, Hen, Blendy, Lucki, & Therapeutics, 2001), the finding that Tph2^-/-^ rats were less anxious may also be due to serotonin-mediated developmental or compensatory changes contribute to the anxiolytic profile.

As 5-HT regulates the aggression in both sexes, enhanced serotonergic activity could inhibit intermale aggression, while hindering 5-HT signaling will stimulate aggression (Carrillo, Ricci, Coppersmith, & Melloni, 2009; Yanowitch & Coccaro, 2011). Serotonin transporter knockout rats exhibit less aggression, more prosocial behaviors with a high sensitivity to social stimuli (Homberg & Lesch, 2011). In our case, Tph2^-/-^ rats had outburst aggressive behaviors almost immediately when housed together with another rat in a novel environment (Supplementary Fig1), as reported Tph2^-/-^ rats have more dense social networks, a more unstable hierarchy and normal social memory (Alonso et al., 2021). Therefore, we propose that Tph2^-/-^ rats have a deficit in updating environmental information, leading to disrupted transmission of social information like hierarchy and social network etc. At the same time, we noticed that Tph2^-/-^ rats spent more time on social contact with their assigned partner, but in a ‘antisocial’ manner with increased mounting behavior. As the animals were tested in male-male social interactions, the mounting behavior might be an act of showing social dominance which is in line with our previous finding in the resident intruder test (Peeters et al., 2019).

The reduced anxiety in Tph2^-/-^ rats may relate to altered oxytocin signaling. Oxytocin infusion into the prelimbic cortex decreased anxiety-like behavior, and pharmacological blockade of the oxytocin receptor prevented this anxiolytic effect, indicating that the anxiolytic effects of oxytocin are mediated, at least in part, through oxytocin receptors in the prelimbic cortex (Sabihi, Dong, Maurer, Post, & Leuner, 2017). Although we did not measure oxytocin levels and oxytocin receptor mRNA expression levels in the same animals, it is well possible that the anxiolytic phenotype of Tph2^-/-^ rats related to elevated oxytocin levels in the medial frontal cortex and enhanced oxytocin receptor expression in the prelimbic cortex. Besides, amygdala plays a key role in emotional processing (LeDoux, 2000) including anxiety, fear learning and memory (Duvarci & Pare, 2014; Janak & Tye, 2015) with γ-aminobutyric acid-ergic (GABAergic) interneurons serving critically for some inhibitory circuits (Stefanits et al., 2018). Presumably, serotonin could alter the GABAergic tone via 5-HT2A receptors (Jiang et al., 2009; McDonald & Mascagni, 2007; Rainnie, 1999). Meanwhile, oxytocin also serves as a potent modulator of inhibitory GABA transmission in the central amygdala. For instance, oxytocin infusion into the central amygdala increased GABA activity in this region (Huber, Veinante, & Stoop, 2005). In line with a previous report that oxytocin infusion into central amygdala could decrease anxiety (Bale, Davis, Auger, Dorsa, & McCarthy, 2001), in our experiment Tph2^-/-^ rats exhibit a lower anxiety level with the oxytocin levels being largely increased in the central nucleus of the amygdala. We therefore suspect that increased oxytocin in this nucleus lowers anxiety levels in Tph2^-/-^ rats by enhancing GABA transmission. The hippocampus can be functionally segmented into dorsal, intermediate and ventral compartments, with the dorsal part mediating cognitive functions and the ventral part implicated in stress, emotion and affect (Dale et al., 2016; Fanselow & Dong, 2010). Previously, it has been reported that a serotoninergic lesion of the ventral hippocampus leads to increased anxiety-like behaviors in the elevated plus maze, showing that serotonin has an anxiety dampening role in the ventral hippocampus (Tu et al., 2014). Surprisingly, in our Tph2^-/-^ rat model, under conditions of life-long deficiency of brain serotonin, rats expressed reduced anxiety. At the same time, we noticed that oxytocin receptor mRNA expression levels were mostly increased in the ventral, but not dorsal compartment of Tph2^-/-^ rats. As intracerebroventricular infusion of oxytocin into the lateral ventricle has anxiolytic effects (Peters, Slattery, Uschold-Schmidt, Reber, & Neumann, 2014; Windle, Shanks, Lightman, & Ingram, 1997), the decreased anxiety as observed in Tph2^-/-^ rats may relate in part to increased oxytocin signaling in the hippocampus. Further investigation is needed to delineate the specific role of oxytocin in the hippocampal subregions and their contribution to Tph2^-/-^ behavior.

Also, the altered social behaviors in Tph2^-/-^ rats may relate to altered oxytocin signaling. The prelimbic cortex participates in the regulation of social interaction (Gonzalez et al., 2000) and oxytocin regulates social approach and preference behaviors (Lukas et al., 2011). Therefore, together with social interaction data from our experiment, we propose that oxytocin in the prelimbic cortex promotes social interaction in Tph2^-/-^ rats. Selective deletion of oxytocin receptors on serotonergic dorsal raphe neurons reduced resident-intruder aggression in males (Pagani et al., 2015). In line with this finding, the oxytocin receptor mRNA expression level in the dorsal raphe nucleus is greatly increased in Tph2^-/-^ rats, which may explain their increased aggressiveness during social interaction. As the change of oxytocin in the dorsal raphe nucleus is slightly decreased in Tph2^-/-^ rats, the increased oxytocin receptor mRNA expression levels could reflect a compensation for reduced oxytocin levels in this region. At the same time, altered GABA transmission in the amygdala also results in exaggerated fear which may explain the high aggressiveness level of Tph2^-/-^ rats during social interaction.

Although in human beings Tph2 complete dysfunction is a very rare situation, there is an association between Tph2 polymorphisms and neuropsychiatric disorders (Zhang, Beaulieu, Gainetdinov, & Caron, 2005; Xiaodong Zhang et al., 2005). Tph2 knockout rat magnifies the phenotype and provides information in the context of serotonin and transdiagnostic behavior. At the same time, some limitations should be taken into account. We only tested male animals, while sex difference could impact the development of oxytocin system (Tamborski, Mintz, & Caldwell, 2016) and oxytocin-dependent behaviors (Dumais, Bredewold, Mayer, & Veenema, 2013). Besides, due to the small brain-punching sample volume, samples used to assess oxytocin levels in the hippocampus and medial prefrontal cortex involve a mixture of subregions.

In conclusion, we demonstrated that rats lacking the Tph2 display a series of behavioral changes which gives us more insights into the effects of long-term serotonin deficiency. Meanwhile, the behavioral changes originating from congenital brain serotonin deficiency sometimes are different from acquired short-term serotonin deficiency due to medical intervention, which suggests that compensatory pathways developed in Tph2^-/-^ rats, with participation of the oxytocin system. The overall increase in oxytocin levels and receptor expression suggests that interventions decreasing oxytocin signaling may have the potential to normalize the anxiolytic and anti-social behavior in those suffering from low Thp2 availability.

## Acknowledgments

The study was supported by the EU H2020 MSCA ITN project “Serotonin and Beyond” (N 953327)

